# Optical Biopsy using a neural network to predict functional state from photos of wounds

**DOI:** 10.1101/2022.09.26.509543

**Authors:** Joe Teague, Damien Socia, Gary An, Stephen Badylak, Scott Johnson, Peng Jiang, Yoram Vodovotz, R. Chase Cockrell

**Affiliations:** Department of Surgery, University of Vermont, 89 Beaumont Ave, Given D319 Burlington, VT 05405; McGowan Institute of Regenerative Medicine, University of Pittsburgh, 450 Technology Drive, Suite 300, Pittsburgh, PA 15219-3110; Cleveland State University, Center for Gene Regulation in Health and Disease (GRHD), 2121 Euclid Avenue, SR 353, Cleveland, OH 44115; Department of Surgery, University of Pittsburgh, W944 Biomedical Sciences Tower, 200 Lothrop St., Pittsburgh, PA 15213

**Keywords:** Machine Learning, Image Processing, Volumetric Muscle Loss, Wound, Soft Tissue Trauma, Gene Expression

## Abstract

**Background:** The clinical characterization of the functional status of active wounds remains a considerable challenge that at present, requires excision of a tissue biopsy. In this pilot study, we use a convolutional Siamese neural network architecture to predict the functional state of a wound using digital photographs of wounds in a canine model of volumetric muscle loss (VML).

**Materials and Methods:** Images of volumetric muscle loss injuries and tissue biopsies were obtained in a canine model of VML. Gene expression profiles for each image were obtained using RNAseq. These profiles were then converted to functional profiles using a manual review of validated gene ontology databases. A Siamese neural network was trained to regress functional profile expression values as a function of the data contained in an extracted image segment showing the surface of a small tissue biopsy. Network performance was assessed in a test set of images using Mean Absolute Percentage Error (MAPE).

**Results:** The network was able to predict the functional expression of a range of functions based with a MAPE ranging from ∼5% to ∼50%, with functions that are most closely associated with the early-state of wound healing to be those best-predicted.

**Conclusions:** These initial results suggest promise for further research regarding this novel use of ML regression on medical images. The regression of functional profiles, as opposed to specific genes, both addresses the challenge of genetic redundancy and gives a deeper insight into the mechanistic configuration of a region of tissue in wounds. As this preliminary study focuses on the first 14 days of wound healing, future work will focus on extending the training data to include longer time periods which would result in additional functions, such as tissue remodeling, having a larger presence in the training data.

## Introduction and Rationale

Mammalian wound healing from volumetric muscle loss (VML) injury is a process that proceeds in a sequence of defined phases characterized by the influx and population dynamics of various cell types, the temporal composition of extracellular matrix and deposited tissue, and the extracellular mediator milieu which comprises a range of effectors including reactive oxygen species, cytokines, proteins, etc. [1-4]. Despite this mechanistic understanding, it is difficult to quantify the internal (e.g., protein concentrations) configuration of the wound and its dynamics without invasive sampling, such as by excising a tissue biopsy, which itself further perturbs the wounds and alters wound healing dynamics [5]. This is particularly relevant when trying to predict the likelihood of successful healing or wound failure [6]. We posit that an effective characterization of the mechanistic state of a wound can both lead to the development of new therapeutics based on fundamental mechanistic knowledge (recognizing that the heterogeneity of a clinical trial population often confounds the results) and to individual clinical benefits through the development of personalized treatment regimens [7, 8].

Though digital photography is frequently used in the clinical setting to establish a record of wound healing progression, the ability to translate those photographs into actionable insights has been limited. We posit that this is partially due to the inability of the clinician to perceive subtle but consistent spatial/color patterning among images, a task at which Artificial Neural Networks (ANNs) excel [9-11]. Machine Learning (ML) in general and ANNs specifically have been applied to a wide range of clinical/medical image analysis tasks ranging from classifying the presence of a disease in an x-ray to outlining specific tumors in medical images [12-16]. In previous work [17], we demonstrated that digital photography could provide both a qualitative and quantitative (for certain mediators that are correlated with wound appearance) record of the temporal evolution of a VML wound in a canine model of volumetric muscle loss. In this work, we utilized a convolutional neural network (CNN), a specific type of ANN, to regress/predict the expression of certain genes in a biopsy-sized volume of tissue, given a photograph of the surface of the biopsy.

The task of gene expression regression/prediction is complicated by the issue of genetic redundancy [18, 19], in which multiple genes can perform the same biochemical function; the consequence of this is that in a genetically heterogenous cohort, distinct individuals can present the same (or a very similar) phenotype while their gene expression profiles are significantly different. We speculate that genetic redundancy is an impediment to the accurate prediction of a wide range of gene expression values; to address this difficulty, we instead propose to regress the functional state of the wound rather than a specific gene expression profile.

## Methods

### Wound Images

As described in [17], high-resolution digital photographs were obtained from a canine model of volumetric muscle loss (VML) injury [20]. The experimental protocol was approved by the University of Pittsburgh Institutional Animal Care and Use Committee as part of a separate project, including the collection of tissue biopsies and the taking of photographs. In brief, dogs were anesthetized. After aseptic preparation of the surgical site the skin was incised to expose the biceps femoris muscle. Approximately 75% of the muscle, including vasculature and nerve, was excised surgically, creating an approximately 10 cm long x 4 cm wide x 2 cm deep VML defect. Hemorrhage was controlled with cautery as needed. Dry Tefla non-adherent gauze dressings were applied and the animals allowed to recover. At various intervals spanning 1, 2, 4, 7, 10 and 14 days after wounding, the animals were re-anesthetized and 6 mm diameter x 18 mm deep biopsies taken with a 6-mm Sklar Tru-Punch disposable biopsy punch from various portions of the wound. Digital photographs were taken of the wounds using a Sony Cybershot 20.4 megapixel camera with no optical or digital zoom prior to each biopsy episode, and locations of the biopsies noted via post-processing marking of the photos. Distance from the wounds (∼ 6 inches) and lighting were kept as consistent as possible for each photograph, with the entire wound captured within a single photograph.

### RNA-Seq

As described in [17], total RNAs were isolated from dog biopsies using trizol (ThermoFisher) and chloroform phase separations followed by the RNeasy mini protocol (Qiagen) with optional on-column DNase digestion (Qiagen). One-hundred nanograms of total RNA was used to prepare sequencing libraries using the LM-Seq (Ligation Mediated Sequencing) protocol [21]. RNAs were selected using the NEB Next Poly A+ Isolation Kit (NEB). Poly A+ fractions were eluted, primed, and fragmented for 7 minutes at 85°C. First-strand cDNA synthesis was performed using SmartScribe Reverse Transcriptase (Takara Bio USA) and RNA is then removed. cDNA fragments were purified with AMpure XP beads (Beckman Coulter). The 5*’* adapter was ligated, and 18 cycles of amplification were performed. These final indexed cDNA libraries were quantified, normalized, multiplexed, and run as single-end reads on the HiSeq 3000 (Illumina). RNA-seq reads were mapped to the dog genome and annotated protein coding genes (version: canFam3) using Bowtie (v0.12.8) [22] allowing up to 2-mismatches. The gene expected read counts and Transcripts Per Million (TPM) were estimated by RSEM (v1.2.3) [23]. This process returned results for 15388 genes.

### Determination of Functional State

In order to determine the functional state of a given biopsy, we first aggregated list of functions associated with each of 2,749 genes that were expressed at a level of greater than 2 TPM from the AmiGO Gene Ontology database [24]. We then cleaned this data by: removing duplicate categories for a single gene, filtering for genes expressed by *canis lupus familiaris*, and removing genes for which no functional annotation is available. Each gene function was then manually reviewed using the UniProt [25], OMIM [26], and GeneCards [27] databases. This resulted in a hierarchical scheme which is illustrated in figure 1; using this method, we have stratified the functions of each gene at three levels (Fig 1, right panel), to allow for different levels of resolution at which the functional regression/prediction can be performed. Specific parent process functions can be seen in the left panel of Fig. 1. In Fig. 2, we show specific examples for the sub-stratification of a parent process (e.g., inflammation) to the first child process (e.g., inflammation inhibition or inflammation regulation). Ultimately, we utilized 485 functions to develop functional profiles. Functional profiles were normalized compositionally, such that the functional expression for an individual biopsy summed to one.

**Figure 1:**
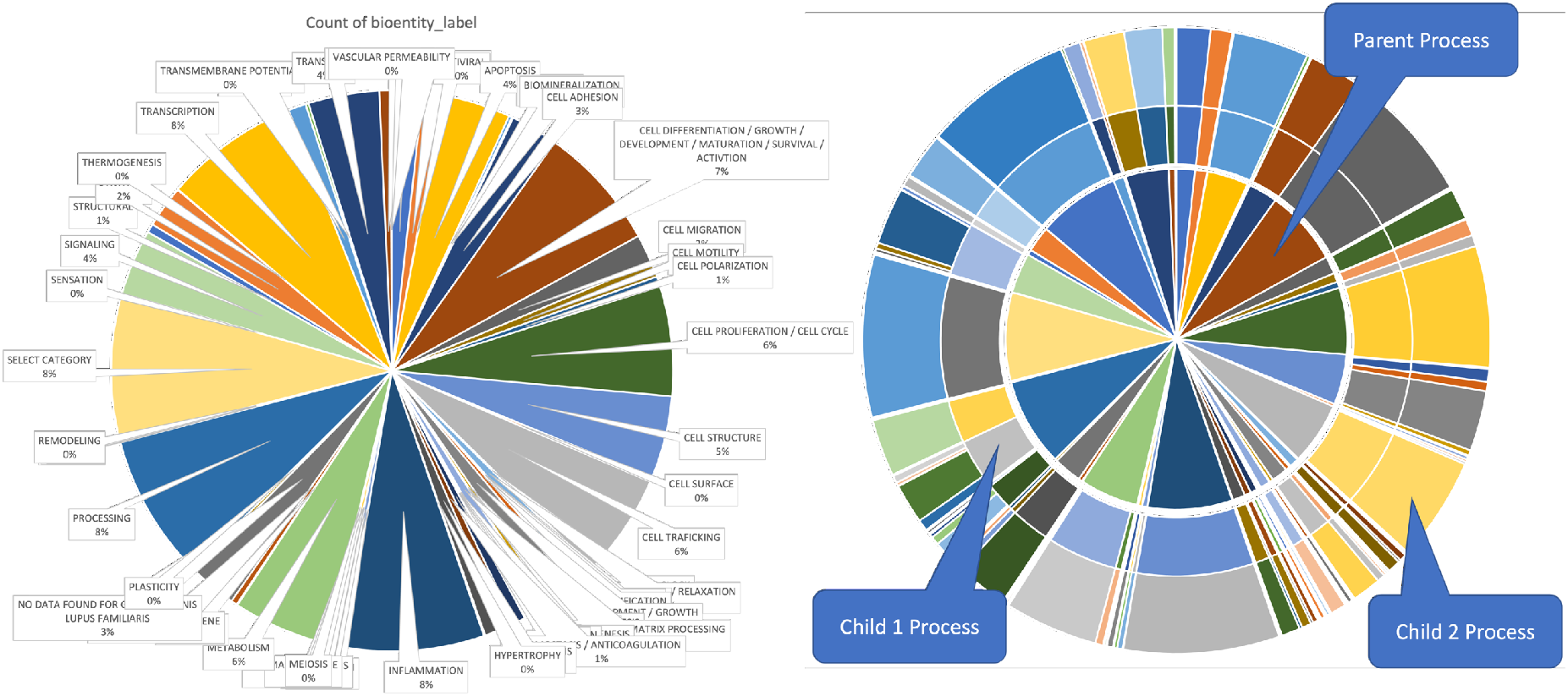
On the left, we present an overview of the distribution of functional parent processes used in this analysis. On the right, we present an illustration of how the parent processes (the center disk) are sub-stratified for further analysis.

**Figure 2:**
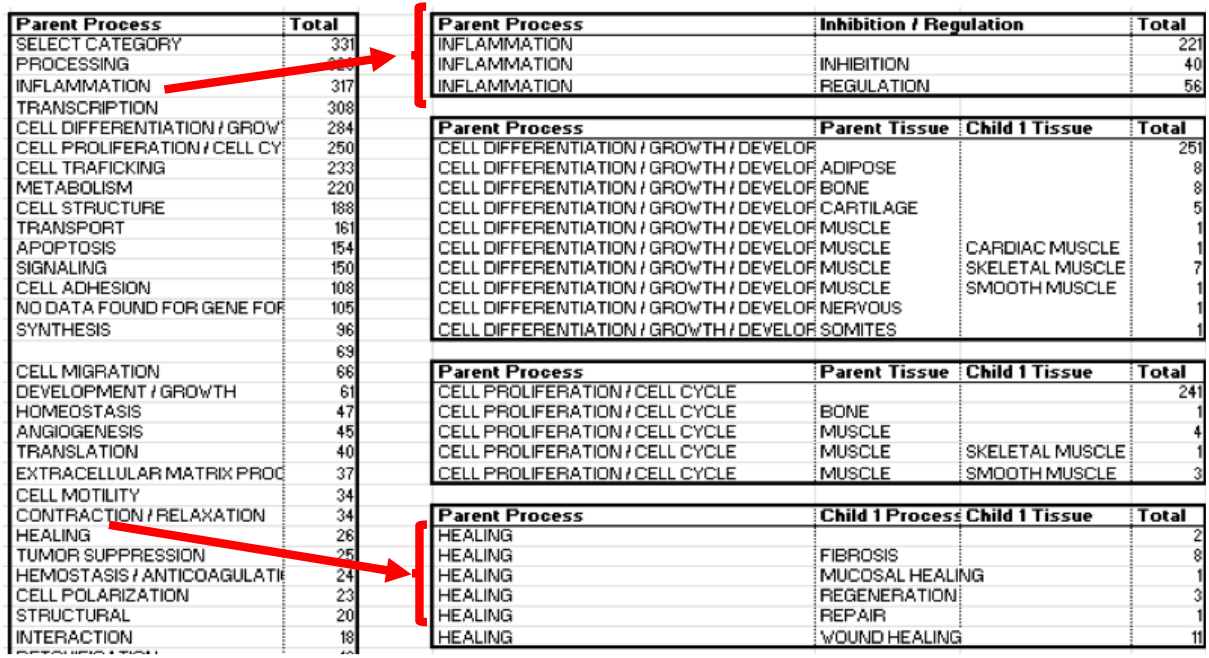
An example of sub-stratification from parent functional process to child 1 functional process to

### Functional State Regression

In order to regress the functional state of the wound, we began with a dataset containing 285 images with corresponding functional profiles (determined from RNAseq, as described above). Each image was a 200 × 200-pixel area which contained the entire surface of the biopsy. To compensate for the small size of our dataset, we utilized three techniques: ensemble networks, data augmentation, and semi-supervised learning.

Ensemble networks [28] reduce the training burden by training multiple (relatively) smaller models rather than a single large network. The function groupings of the genes have a hierarchical structure to them with the highest level being the broadest category and the lowest level being the most specific. Individual NNs were then trained for each high-level functional group of interest. The outputs for each NN were the lowest level functional group in that category.

Data augmentation was used to add variation to each image as the network training proceeds. Each time an image was selected for training, there was a random chance that it could be mirrored across either central vertical or horizontal planes [29], or it could be randomly rotated up to 10 degrees [30]. These augmentations ameliorate the dangers of overfitting a small data set through correlating multiple image presentations with the same functional profile expression values.

Semi-supervised learning, [31, 32] is a machine learning method applicable to datasets that have both labeled and unlabeled data. This technique makes an excellent augmentation to our data as biopsy images are extracted from high-resolution images of the entire wound, thus there is ample unlabeled data for each labeled data sample. Using this technique, random 200 × 200-pixel images sections were obtained from each wound, resulting in an additional 4156 unlabeled biopsy-sized images for training. Each of these was manually reviewed prior to training to ensure that they contained only the wound (i.e., there was no skin or fur visible in the biopsy-sized images). We note that while this unlabeled data is not perfect/ideal, it does improve the ultimate performance of the trained networks.

We utilized a metric-based semi-supervised training algorithm [33] to train the functional regression model on the partially labeled dataset. We used a dual-component training scheme, in which the first component trained the image feature space on the partially unlabeled dataset and the second component trained the regressor on that feature space using only the labeled data. To train the feature space, we used a Siamese network architecture [34, 35], which is a network architecture that utilized two identical and parallel networks, allowing for the network to train on two images simultaneously. The output of this is then two feature spaces that are compared using a contrastive loss function [36], which determines the difference between the features space and trains the model to output similar feature spaces for similar images and dissimilar feature spaces for dissimilar images. To determine the similarity between images, all the images were paired and either the difference between functional profile expression or the Euclidean distance in the event that one of the images was unlabeled. The top 5% of labeled and unlabeled pairs were then considered to be similar. We note that as the labeled pairs contain more information so they were weighted more heavily in training.

The Siamese network comprised only convolutional layers, with Resnet34 giving the best results. For the second stage of training the convolutional layers trained in the first stage were frozen and a linear network with one hidden layer was trained. The linear layers take the feature space from the first model and regresses the gene functional groups. Together the two models run in series are able to regress the gene functional groups from an image of a wound.

## Results

Model performance was evaluated using the Mean Absolute Percent Error (MAPE). MAPE is a metric that has been used to measure prediction accuracy for forecasting and regression [37]. The formula for MAPE is:

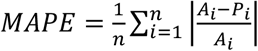

where *n* represents the number or fitted points, *A*_&_ represents the actual value, and *P*_&_ represents the predicted value. We note that while the minimum value for this error is 0 (the prediction is perfect), the maximum value is unbounded, and consequently, can be very high for genes or functions that are expressed at very low levels. This constraint leads to an unreliability in the prediction of Child 2 functional processes – specifically, the number of processes is high so each one makes up only a tiny fraction of the total functional profile. Conversely, when considering only parent processes, the level of resolution is too granular to make any intelligent or actionable predictions (e.g., an exemplar parent process is *‘*Inflammation [see Fig. 2], which itself it not informative). In Fig. 3, we present results for the top 11 predicted functions with associated 95% confidence intervals. In table 1, we present the MAPE and mean functional expression levels (parts per million) for all functions that were predicted with an error <50%.

**Figure 3:**
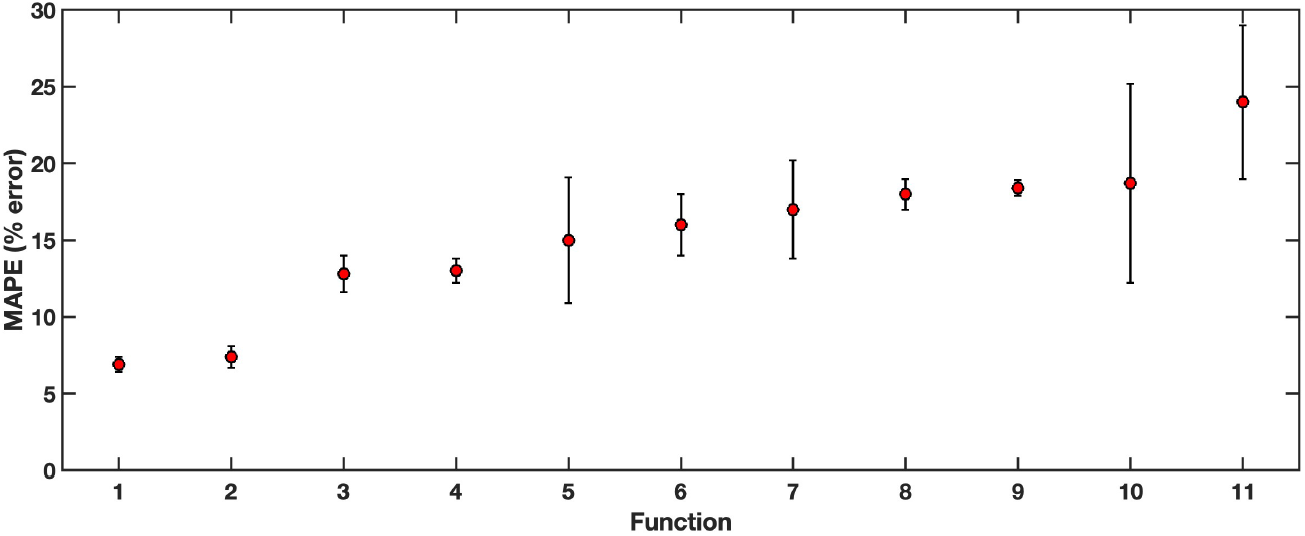
Mean average percent errors for functional regression are displayed for: 1. Cell Proliferation (General), 2. Cell Trafficking (General), 3. Cell Migration (General), 4. Cell Proliferation Inhibition (General) 5. Growth Inhibition of Bone, 6. Growth of Neurite, 7. Differentiation/Activation of Neutrophils, 8. Growth regulation of nervous system,9. Differentiation/Activation of WBC*’*s,10. Development/Activation of Neurons,11. Development/Growth of Nerve

**Table 1:** Child 2 functional processes, Mean Average Percent Error, and total functional expression (parts per million).

| Function | MAPE | Mean Expression |
| --- | --- | --- |
| CELL PROLIFERATION / CELL CYCLE REGULATION | 0.06921429 | 15568.2447 |
| CELL TRAFICKING | 0.07438151 | 39120.1598 |
| CELL MIGRATION | 0.12791456 | 7529.1112 |
| CELL PROLIFERATION / CELL CYCLE | 0.13497846 | 13796.962 |
| CELL PROLIFERATION / CELL CYCLE INHIBITION | 0.14326078 | 10493.8452 |
| DEVELOPMENT / GROWTH INHIBITION OF BONE | 0.15076176 | 465.333168 |
| DEVELOPMENT / GROWTH OF NEURITE | 0.16421275 | 352.193562 |
| ('CELL DIFFERENTIATION / GROWTH / DEVELOPMENT / MATURATION / SURVIVAL / ACTIVTION OF HEMATOPOIETIC CELLS'.) | 0.17568658 | 1984.14829 |
| ('CELL DIFFERENTIATION / GROWTH / DEVELOPMENT / MATURATION / SURVIVAL / ACTIVTION OF NEUTROPHILS'.) | 0.17990649 | 1189.46153 |
| DEVELOPMENT / GROWTH REGULATION OF NERVOUS SYSTEM | 0.18068103 | 186.181147 |
| ('CELL DIFFERENTIATION / GROWTH / DEVELOPMENT / MATURATION / SURVIVAL / ACTIVTION OF WBCs'.) | 0.18437668 | 3203.8455 |
| ('CELL DIFFERENTIATION / GROWTH / DEVELOPMENT / MATURATION / SURVIVAL / ACTIVTION OF NEURONS'.) | 0.18773514 | 3088.61104 |
| ('CELL DIFFERENTIATION / GROWTH / DEVELOPMENT / MATURATION / SURVIVAL / ACTIVTION'.) | 0.19998091 | 8394.51879 |
| ('CELL DIFFERENTIATION / GROWTH / DEVELOPMENT / MATURATION / SURVIVAL / ACTIVTION REGULATION OF HEMATOPOIETIC CELLS'.) | 0.21381868 | 334.457387 |
| CELL MIGRATION REGULATION | 0.21790823 | 1249.22842 |
| CELL MIGRATION OF ENDOTHELIAL CELLS | 0.23329506 | 339.46842 |
| ('CELL DIFFERENTIATION / GROWTH / DEVELOPMENT / MATURATION / SURVIVAL / ACTIVTION OF EPIDERMAL CELLS'.) | 0.24162359 | 422.938659 |
| CELL MOTILITY | 0.24430771 | 11105.3684 |
| ('CELL DIFFERENTIATION / GROWTH / DEVELOPMENT / MATURATION / SURVIVAL / ACTIVTION OF ERYTHROCYTES'.) | 0.24557471 | 799.122943 |
| DEVELOPMENT / GROWTH OF NERVE | 0.24705462 | 528.37423 |
| CELL MIGRATION OF SMOOTH MUSCLE CELLS | 0.26314071 | 645.359256 |
| CELL MOTILITY REGULATION | 0.27588635 | 6156.79707 |
| ('CELL DIFFERENTIATION / GROWTH / DEVELOPMENT / MATURATION / SURVIVAL / ACTIVTION OF KERATINOCYTES'.) | 0.27754401 | 1938.01672 |
| CELL MIGRATION OF T CELLS | 0.28173003 | 306.78646 |
| ('CELL DIFFERENTIATION / GROWTH / DEVELOPMENT / MATURATION / SURVIVAL / ACTIVTION OF ADIPOCYTES'.) | 0.28228317 | 1319.922 |
| SECRETION REGULATION OF SYNAPSE | 0.28346258 | 234.382479 |
| ('CELL DIFFERENTIATION / GROWTH / DEVELOPMENT / MATURATION / SURVIVAL / ACTIVTION OF B CELLS'.) | 0.28358586 | 1348.54189 |
| STRUCTURAL | 0.28688075 | 132.797145 |
| DEVELOPMENT / GROWTH OF NERVOUS SYSTEM | 0.29746975 | 2628.99185 |
| CELL PROLIFERATION / CELL CYCLE OF SMOOTH MUSCLE CELLS | 0.29835947 | 645.359256 |
| CELL TRAFICKING OF RIBOSOMES | 0.31518554 | 571.519001 |
| ('CELL DIFFERENTIATION / GROWTH / DEVELOPMENT / MATURATION / SURVIVAL / ACTIVTION INHIBITION'.) | 0.31734605 | 392.498488 |
| DEVELOPMENT / GROWTH INHIBITION OF NEURITE | 0.32630352 | 717.793173 |
| CELL PROLIFERATION / CELL CYCLE OF NEURONS | 0.32753865 | 375.247005 |
| DEVELOPMENT / GROWTH OF NEURONS | 0.34861897 | 150.887649 |
| CELL TRAFICKING OF HORMONES | 0.35776058 | 255.419273 |
| ('CELL DIFFERENTIATION / GROWTH / DEVELOPMENT / MATURATION / SURVIVAL / ACTIVTION OF OSTEOBLASTS'.) | 0.36539425 | 549.394796 |
| CELL TRAFICKING OF NEURONS | 0.37114547 | 97.1478886 |
| ('CELL DIFFERENTIATION / GROWTH / DEVELOPMENT / MATURATION / SURVIVAL / ACTIVTION OF T CELLS'.) | 0.37950688 | 3947.05823 |
| ('CELL PROLIFERATION / CELL CYCLE TRANSCRIPTION'.) | 0.38043112 | 118.663123 |
| CELL PROLIFERATION / CELL CYCLE REGULATION OF GLOMERULAR CELLS | 0.38408548 | 141.626878 |
| CELL PROLIFERATION / CELL CYCLE OF KERATINOCYTES | 0.39477446 | 203.524481 |
| ('CELL DIFFERENTIATION / GROWTH / DEVELOPMENT / MATURATION / SURVIVAL / ACTIVTION OF EPITHELIAL CELLS'.) | 0.39589214 | 157.504169 |
| DEVELOPMENT / GROWTH OF BLOOD VESSELS | 0.40265949 | 643.36836 |
| DEVELOPMENT / GROWTH OF LYMPHATIC VESSELS | 0.40285128 | 643.36836 |
| ('CELL DIFFERENTIATION / GROWTH / DEVELOPMENT / MATURATION / SURVIVAL / ACTIVTION INHIBITION OF WBCs'.) | 0.40394941 | 183.213134 |
| CELL TRAFICKING OF NEUROTRANSMITTERS | 0.41131179 | 255.419273 |
| CELL PROLIFERATION / CELL CYCLE OF T CELLS | 0.41845448 | 651.586849 |
| ('CELL DIFFERENTIATION / GROWTH / DEVELOPMENT / MATURATION / SURVIVAL / ACTIVTION REGULATION'.) | 0.44578599 | 20498.1573 |
| ('CELL DIFFERENTIATION / GROWTH / DEVELOPMENT / MATURATION / SURVIVAL / ACTIVTION REGULATION OF T CELLS'.) | 0.45379946 | 211.165867 |
| CELL MIGRATION OF FIBROBLASTS | 0.45898248 | 174.750242 |
| GROWTH GUIDANCE OF NEURITE | 0.46443711 | 462.373407 |
| ('ESTROGEN MEDIATED CELL DIFFERENTIATION / GROWTH / DEVELOPMENT / MATURATION / SURVIVAL / ACTIVTION OF MACROPHAGES'.) | 0.47237411 | 802.476697 |

## Discussion

While the use of ML for clinical/medical applications is increasingly common, these techniques are typically used for classification [38-42] or image segmentation [43, 44]. This work attempts to go beyond that by regressing a full functional profile for each tissue biopsy. There is some related work in cancer [45] that attempts to infer gene expression through the identification of particular markers for tumor behavior; however, we note a fundamental distinction between this and that work is that this requires a tissue biopsy (with the ultimate goal of replacing the tissue biopsy).

We note that this work has major limitations, the first of which is the paucity of training data. While we attempt to address this limitation with various data augmentation techniques, none of these would be as good as including additional data. While the above-described models have found generalizable patterns in the dataset used, it is difficult to determine the degree of generalizability, i.e., how these results would change with the inclusion of more dogs or different breeds of dogs. Additionally, we note that the biopsy images were sampled form a canine VML model within 14 days post-injury, and as such, many of the remodeling functions are not present or minimally present (e.g., the formation of fibrotic scar) in the training data. Lastly, all of the wounds utilized in this study appeared to be on a trajectory to heal successfully (with fibrotic scar) at the end of the experiment, thus, functions associated with wound failure were not present in this training data.

Given the extreme heterogeneity in gene expression, both between dogs and within a single wound, many functions were predicted with a relatively accurate degree of fidelity. In general, the most accurate predictions were those of the general class, meaning that they apply to all cell types more or less equally. However, certain cell types had functional predictions significantly more accurate than others; these included both neurons and inflammatory cells such as neutrophils or white blood cells.

The regression of functional profiles as opposed to specific genes [17] greatly increased the depth of wound characterization that is tractable to obtain non-invasively. Despite the limitations of this study, we believe that it shows promise as a technique to gain functional/mechanistic insight into the state of wounds in a real-time (as often as a photograph can be taken) and non-invasive manner. In future work, we will expand the training data, both in terms of number of images utilized as well as the timing of image acquisition post wound such that later-term functional processes are included.

## References

1. Stroncek JD, Reichert WM. Overview of wound healing in different tissue types. Indwelling neural implants: strategies for contending with the in vivo environment. 2008;1:3–41.

2. McCarty SM, Percival SL. Proteases and delayed wound healing. Advances in wound care. 2013;2(8):438–47.

3. Martin P, Leibovich SJ. Inflammatory cells during wound repair: the good, the bad and the ugly. Trends Cell Biol. 2005;15(11):599–607.

4. Witte MB, Barbul A. General principles of wound healing. Surg Clin North Am. 1997;77(3):509–28.

5. Amler MH. Disturbed healing of extraction wounds. J Oral Implantol. 1999;25(3):179–84.

6. Forsberg JA, Potter BK, Polfer EM, Safford SD, Elster EA. Do inflammatory markers portend heterotopic ossification and wound failure in combat wounds? Clinical Orthopaedics and Related Research®. 2014;472(9):2845–54.

7. Kassab GS, An G, Sander EA, Miga MI, Guccione JM, Ji S, et al. Augmenting surgery via multi-scale modeling and translational systems biology in the era of precision medicine: a multidisciplinary perspective. Annals of biomedical engineering. 2016;44(9):2611–25.

8. Ziraldo C, Solovyev A, Allegretti A, Krishnan S, Henzel MK, Sowa GA, et al. A computational, tissue-realistic model of pressure ulcer formation in individuals with spinal cord injury. PLoS computational biology. 2015;11(6):e1004309.

9. Egmont-Petersen M, de Ridder D, Handels H. Image processing with neural networks—a review. Pattern recognition. 2002;35(10):2279–301.

10. He K, Zhang X, Ren S, Sun J, editors. Deep residual learning for image recognition. Proceedings of the IEEE conference on computer vision and pattern recognition; 2016.

11. Wu Z, Shen C, Van Den Hengel A. Wider or deeper: Revisiting the resnet model for visual recognition. Pattern Recognition. 2019;90:119–33.

12. Lo S-CB, Chan H-P, Lin J-S, Li H, Freedman MT, Mun SK. Artificial convolution neural network for medical image pattern recognition. Neural networks. 1995;8(7-8):1201–14.

13. Jiang J, Trundle P, Ren J. Medical image analysis with artificial neural networks. Computerized Medical Imaging and Graphics. 2010;34(8):617–31.

14. Lakhani P, Sundaram B. Deep learning at chest radiography: automated classification of pulmonary tuberculosis by using convolutional neural networks. Radiology. 2017;284(2):574–82.

15. Antony J, McGuinness K, O’Connor NE, Moran K, editors. Quantifying radiographic knee osteoarthritis severity using deep convolutional neural networks. 2016 23rd International Conference on Pattern Recognition (ICPR); 2016: IEEE.

16. Rajkomar A, Lingam S, Taylor AG, Blum M, Mongan J. High-throughput classification of radiographs using deep convolutional neural networks. Journal of digital imaging. 2017;30(1):95–101.

17. Schumaker G, Becker A, An G, Badylak S, Johnson S, Jiang P, et al. Optical Biopsy Using a Neural Network to Predict Gene Expression From Photos of Wounds. J Surg Res. 2022;270:547–54.

18. Láruson ÁJ, Yeaman S, Lotterhos KE. The importance of genetic redundancy in evolution. Trends Ecol Evol. 2020;35(9):809–22.

19. Brookfield JF. Genetic redundancy. Adv Genet. 1997;36(C):137–55.

20. Turner NJ, Badylak JS, Weber DJ, Badylak SF. Biologic scaffold remodeling in a dog model of complex musculoskeletal injury. Journal of Surgical Research. 2012;176(2):490–502.

21. Hou Z, Jiang P, Swanson SA, Elwell AL, Nguyen BK, Bolin JM, et al. A cost-effective RNA sequencing protocol for large-scale gene expression studies. Sci Rep. 2015;5:9570. doi: 10.1038/srep09570. PubMed PMID: 25831155; PubMed Central PMCID: PMC4381617.

22. Langmead B, Trapnell C, Pop M, Salzberg SL. Ultrafast and memory-efficient alignment of short DNA sequences to the human genome. Genome Biol. 2009;10(3):R25. doi: 10.1186/gb-2009-10-3-r25. PubMed PMID: 19261174; PubMed Central PMCID: PMC2690996.

23. Li B, Dewey CN. RSEM: accurate transcript quantification from RNA-Seq data with or without a reference genome. BMC Bioinformatics. 2011;12:323. doi: 10.1186/1471-2105-12-323. PubMed PMID: 21816040; PubMed Central PMCID: PMC3163565.

24. Carbon S, Ireland A, Mungall CJ, Shu S, Marshall B, Lewis S, et al. AmiGO: online access to ontology and annotation data. Bioinformatics. 2009;25(2):288–9.

25. UniProt: the universal protein knowledgebase in 2021. Nucleic Acids Res. 2021;49(D1):D480–D9.

26. Amberger JS, Bocchini CA, Schiettecatte F, Scott AF, Hamosh A. OMIM. org: Online Mendelian Inheritance in Man (OMIM®), an online catalog of human genes and genetic disorders. Nucleic Acids Res. 2015;43(D1):D789–D98.

27. Safran M, Dalah I, Alexander J, Rosen N, Iny Stein T, Shmoish M, et al. GeneCards Version 3: the human gene integrator. Database. 2010;2010.

28. Perrone MP, Cooper LN. When networks disagree: Ensemble methods for hybrid neural networks. Brown Univ Providence Ri Inst for Brain and Neural Systems, 1992.

29. Shijie J, Ping W, Peiyi J, Siping H, editors. Research on data augmentation for image classification based on convolution neural networks. 2017 Chinese automation congress (CAC); 2017: IEEE.

30. Quiroga F, Ronchetti F, Lanzarini L, Bariviera AF, editors. Revisiting data augmentation for rotational invariance in convolutional neural networks. International Conference on Modelling and Simulation in Management Sciences; 2018: Springer.

31. Zhu X, Goldberg AB. Introduction to semi-supervised learning. Synthesis lectures on artificial intelligence and machine learning. 2009;3(1):1–130.

32. Zhou X, Belkin M. Semi-supervised learning. Academic Press Library in Signal Processing. 1: Elsevier; 2014. p. 1239–69.

33. Zhu XJ. Semi-supervised learning literature survey. 2005.

34. Melekhov I, Kannala J, Rahtu E, editors. Siamese network features for image matching. 2016 23rd international conference on pattern recognition (ICPR); 2016: IEEE.

35. Gleize M, Shnarch E, Choshen L, Dankin L, Moshkowich G, Aharonov R, et al. Are you convinced? choosing the more convincing evidence with a Siamese network. arXiv preprint 190708971. 2019.

36. Wang F, Liu H, editors. Understanding the behaviour of contrastive loss. Proceedings of the IEEE/CVF conference on computer vision and pattern recognition; 2021.

37. De Myttenaere A, Golden B, Le Grand B, Rossi F. Mean absolute percentage error for regression models. Neurocomputing. 2016;192:38–48.

38. Rostami B, Anisuzzaman D, Wang C, Gopalakrishnan S, Niezgoda J, Yu Z. Multiclass Wound Image Classification using an Ensemble Deep CNN-based Classifier. arXiv preprint 201009593. 2020.

39. Jiang Z, Ardywibowo R, Samereh A, Evans HL, Lober WB, Chang X, et al. A roadmap for automatic surgical site infection detection and evaluation using user-generated incision images. Surgical infections. 2019;20(7):555–65.

40. Hsu J-T, Chen Y-W, Ho T-W, Tai H-C, Wu J-M, Sun H-Y, et al. Chronic wound assessment and infection detection method. BMC medical informatics and decision making. 2019;19(1):1–20.

41. Shenoy VN, Foster E, Aalami L, Majeed B, Aalami O, editors. Deepwound: Automated postoperative wound assessment and surgical site surveillance through convolutional neural networks. 2018 IEEE International Conference on Bioinformatics and Biomedicine (BIBM); 2018: IEEE.

42. Wu J-M, Tsai C-J, Ho T-W, Lai F, Tai H-C, Lin M-T. A Unified Framework for Automatic Detection of Wound Infection with Artificial Intelligence. Applied Sciences. 2020;10(15):5353.

43. Wang C, Anisuzzaman D, Williamson V, Dhar MK, Rostami B, Niezgoda J, et al. Fully automatic wound segmentation with deep convolutional neural networks. Scientific Reports. 2020;10(1):1–9.

44. Li F, Wang C, Liu X, Peng Y, Jin S. A composite model of wound segmentation based on traditional methods and deep neural networks. Computational intelligence and neuroscience. 2018;2018.

45. Schmauch B, Romagnoni A, Pronier E, Saillard C, Maillé P, Calderaro J, et al. A deep learning model to predict RNA-Seq expression of tumours from whole slide images. Nature communications. 2020;11(1):1–15.

